# Continental species distribution and biodiversity predictions depend on modeling grain

**DOI:** 10.1101/2023.10.25.564051

**Authors:** Jeremy M. Cohen, Walter Jetz

## Abstract

As global change accelerates, accurate predictions of species distributions and biodiversity patterns are critical to prevent population declines and biodiversity loss. However, at continental and global scales, these predictions are often derived from species distribution models (SDMs) fit at coarse spatial grains uninformed by ecological processes. Coarse-grain models may systematically bias predictions of distributions and biodiversity if they are consistently over- or under-estimating area with suitable habitat, and this bias may intensify in regions with heterogenous landscapes or with poor data coverage. To test this, we fit presence-absence SDMs characterizing both the summer and winter distributions of 572 North American bird species – nearly the entire avian diversity of the US and Canada – across five spatial grains from 1 to 50 km, using observations from the eBird citizen science initiative. We find that across both seasons, models fit at 1 km performed better under cross-validation than those at coarser scales and more accurately predicted species’ presences and absences at local sites. Coarser-grain models, including models fit at 3 km, consistently under-predicted range area relative to 1 km models, suggesting that coarse-grain estimates of distributions could be missing important habitat. This bias intensified during summer (83% of species) when many birds have smaller ‘operational scales’ via localized home ranges and greater habitat specificity while breeding. Biases were greatest in heterogenous desert and scrubland regions and lowest in more homogenous boreal forest and taiga-dominated regions. When aggregating distributions to produce continental biodiversity predictions, coarse-grain models overpredicted diversity in the west and underpredicted it in the great plains, prairie pothole region and boreal/taiga zones. The modern availability of high-performance computing and high-resolution observational and environmental data provides opportunities to improve continental predictions of species distributions and biodiversity.

## Introduction

Global biodiversity loss is accelerating as changes to climate and land-use alter relationships between species and the environment (Abrahms et al., 2017; Buckley and Jetz, 2008; Pacifici et al., 2015), and researchers, conservationists, and managers rely on accurate predictions of species distributions and richness to assess population declines and biodiversity loss driven by global change (Araújo et al., 2019; Jetz et al., 2008). Although spatial grain size (i.e., sampling resolution) has long been recognized by ecologists as critical to understanding species distributions and biodiversity (Levin, 1992; Wiens, 1989), it is often an afterthought when characterizing species’ environmental niches, predicting species distributions, and estimating richness (Lu and Jetz, 2023). However, models conducted at coarse spatial grains may be compromised because they summarize environmental data over much larger units than areas in which observations are made, creating a spatial mismatch between environmental information and observations (Austin and Van Niel, 2011; Connor et al., 2018). As such, these models may be subject to numerous significant biases affecting the accuracy of species distributions and biodiversity estimates, including the misestimation of range boundaries, range sizes and local biodiversity (Hurlbert and Jetz, 2007; Mertes and Jetz, 2018; Seo et al., 2009). With most current estimates of continental distributions and biodiversity patterns based on coarse-grain models (e.g., Distler et al., 2015; Jung et al., 2021; Mi et al., 2023; Stewart et al., 2022), it remains unknown how these biases are collectively influencing our picture of changing species distributions and biodiversity patterns (Araújo et al., 2019; Thuiller et al., 2019).

Coarse-grain models may be failing to correctly pinpoint utilized habitat or identify important biodiverse areas at local scales for several reasons (A. Lee-Yaw et al., 2022). First, environmental conditions are averaged at coarse grains, leading models to associate species’ occurrences with conditions that do not match the area where the observation was made and truncating niche volume towards environmental means (Lu and Jetz, 2023). Second, coarse-scale models may overestimate the distributions of evenly-dispersed species compared with those with clustered individuals due to the occupancy-scale relationship (He and Condit, 2007). Third, opportunistic, point-based observational data (e.g., GBIF or eBird data) are increasingly utilized to generate large-scale predictions of species distributions and biodiversity (Gaiji et al., 2013; Johnston et al., 2019), replacing atlas data based on systematic surveys of grid cells. However, unevenly distributed spatial gaps between observation points preclude point-based observational data from being useful for estimating distributions or richness over coarse cells covering large areas, especially in poorly sampled regions (e.g., interior Western US, northern Canada).

Several predictions can be generated about the biases inherent in coarse-grain SDMs (Fig. 1). Distributional predictions in heterogenous landscapes (e.g., the American west or prairie pothole region of the northern Great Plains), especially when generated using point-based data, may over- or under-estimate area of occurrence (Oldfather et al., 2020) by designating either all or none of a large cell as suitable for a given species despite only a subset being truly suitable (e.g., an isolated, forested mountain surrounded by lowland desert), resulting in low accuracy of estimated species presence or absence (Fig. 1). In contrast, coarse-grain SDMs may perform and predict similarly to fine-grain SDMs in homogenous landscapes, where environmental conditions are consistent regardless of cell size (Fig. 1). Systematic misestimation of area of occurrence across species can result in artificially high or low biodiversity estimates when distributions are aggregated, a bias that may intensify in regions with heterogenous landscapes or poor data coverage. Such biases are especially concerning because heterogenous landscapes often contain range-limited and habitat-specialist species that are most at risk of declines and extirpations (Clavel et al., 2011). Despite these biases, coarse modeling grains are often chosen based on computational availability (Moudrý and Šímová, 2012) or available environmental data rather than the grain-relevant ecological processes (Mertes and Jetz, 2018), and fine-scale SDMs across numerous species at continental extents have been difficult to produce as they require large amounts of computational power and high-resolution environmental predictors (Beery et al., 2021). However, the recent explosion in the availability of high-performance computing clusters, high-resolution species occurrence data, and remote sensing products summarized at 1 km resolution or below can facilitate the production of fine-scale estimates of species distributions for an increasing number of species (He et al., 2015; Urbina-Cardona et al., 2019).

**Figure 1.**
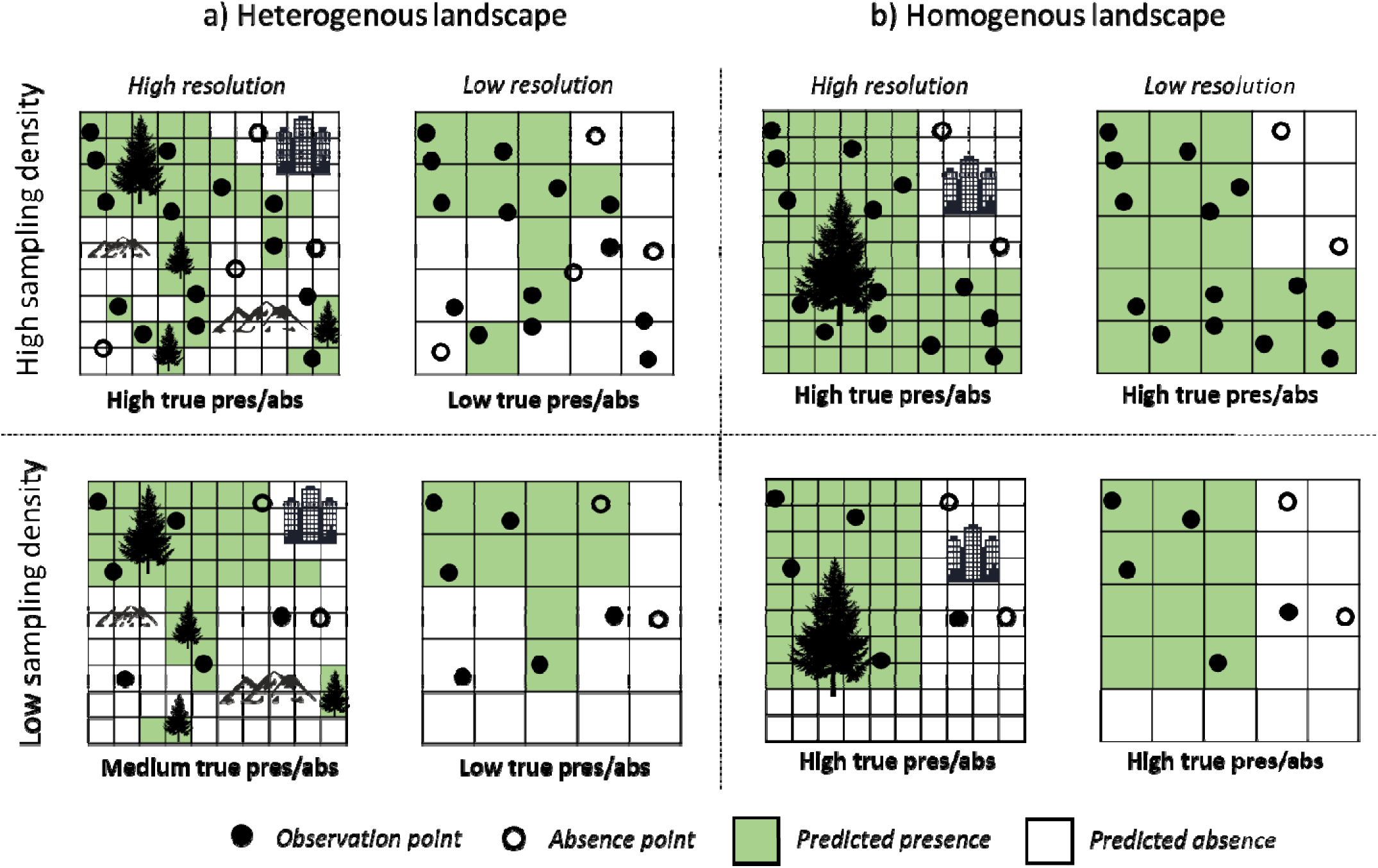
Modeling resolution in heterogenous and homogenous landscapes. (a) Conceptual diagram illustrating that in heterogenous landscapes, small modeling grain (high resolution) of environmental features is likely to result in precise predictions of species presence (green cells) or absence (white cells) across habitat types, as observation points (solid points) and absence points (open points) are well-matched to corresponding environmental features. Predictions are likely to be especially precise when sampling density is high. b) In homogenous landscapes, predictions of species presence or absence may be accurate regardless of grain size, as habitat varies little at small scales. Predictions may also remain accurate even when sampling density is low.

Both simulation studies and empirical, regional-scale work have examined the influence of spatial grain on predicting species distributions (Connor et al., 2018; Gottschalk et al., 2011; Guisan et al., 2007; Seo et al., 2009). However, the implications of grain size on the distribution characterizations or continental biodiversity patterns for a diverse taxon remain underexplored (but see Carroll et al., 2022; Chauvier et al., 2022). Further, it remains unknown whether grain-related bias in predicting species distributions and biodiversity is more prominent during certain portions of the year (Mateo Sánchez et al., 2014), given the tighter home ranges and specific habitat requirements of many species during the breeding season (Zuckerberg et al., 2016). As such, existing coarse-grain models may be misleading applied conservation efforts and projections of biodiversity loss under global change (Jetz et al., 2008; Lecours et al., 2015; Mertes and Jetz, 2018). Estimating continental biodiversity using finer-scale models is crucial to gaining a more accurate picture of biodiversity patterns and properly managing ongoing range shifts and biodiversity loss from regional to local scales (Barbet-Massin and Jetz, 2015; Gillingham et al., 2012; Lecours et al., 2015; Mertes and Jetz, 2018).

Our overarching goal was to determine how the choice of spatial grain influences species-level area of occurrence estimates and aggregated, continental biodiversity metrics across an entire clade and multiple seasons. To accomplish our goal, we independently modeled the seasonal, continental distributions of 572 North American birds – nearly the full diversity of US- and Canada-breeding species – at 1, 3, 5, 10 and 50 km (cell edge lengths). Although few estimates of distributions and diversity are generated at 50 km, we include it as a benchmark to compare against other coarse grains. At each of these five spatial grains, we applied a machine learning algorithm to fit SDMs to tens of millions of georeferenced bird observations obtained via the eBird citizen science initiative (Sullivan et al., 2014), each annotated with a suite of spatial climate, topographic, and landcover covariates upscaled to the appropriate grain size, and predicted thresholded range boundaries for each SDM using a prediction surface with identical resolution. We independently modeled both the summer (breeding) and winter distributions of each species across spatial grains. In total, we fit 4,955 SDMs, one for each combination of species, season, and spatial grain, then aggregated the species-level predictions within each modeling grain and season to develop seasonal, continental estimates of richness and range-size rarity for each grain size. Finally, we evaluated each model via cross-validation with independent testing data and evaluated our aggregate predictions at each spatial grain by comparing local bird checklists at 322 sites with predicted species presences and absences at each site based on our models.

Our motivating questions were as follows: 1) Which choice of spatial grain produces the most accurate predictions of species distributions and biodiversity? 2) How does spatial grain choice alter species-level area-of-occurrence and biodiversity estimates? 3) Which species and regions are prone to demonstrate the most extreme bias at coarse grain? 4) Are grain effects seasonal, perhaps because species are more habitat-specific during summer, when they are breeding? We predicted that coarse-grain models largely misestimate both area-of-occurrence and biodiversity, particularly during summer, and that this bias is more extreme among restricted-range and habitat-specialist species and in regions with heterogenous landscapes.

## Methods

### Spatial modeling extent and species selection

We modeled species distributions across the North American continent – a box with dimensions (179.99°W, 42.68°W, 10.53°S, and 87.11°N), containing Canada, the US, and most of Central America. Although we only modeled species native to the US and Canada and only present biodiversity estimates within this region, modeling the full continental range of each species was necessary to ensure accurate predictions.

We compiled a list of 672 native bird species that annually breed or overwinter in the United States and Canada based on American Birding Association birding codes (2008) updated to the Clements bird taxonomy as of 2021 (Clements, 2007). These codes are a widely-accepted authority to distinguish regularly occurring species (code 1 or 2 species, which we selected) from irregularly occurring vagrants (code 3+). Some species from our original list were eventually excluded because their ranges within our study extent are almost exclusively marine, there were insufficient (<15) data points available, or we were unable to model the species at every spatial grain. We also excluded Hawaiian endemics because most species were restricted to islands smaller than some of our spatial grain sizes. In the end, we modeled 572 species during summer, which is the breeding season for most species, and 419 during winter, as species which overwinter entirely south of the US were excluded.

### Data acquisition

We compiled observational data from eBird, a global citizen science initiative in which users submit checklists containing bird observations that has become widely used for understanding species distributions at high resolution (Sullivan et al., 2014). Users have the option to indicate whether all observed species were recorded on the checklist (“complete” checklists), allowing the inference of absences and presence-absence modelling. Further, users indicate the level of effort involved in each observation by providing the distance traveled, amount of time birding and number of observers (hereafter, effort indicators).

All data compilation, analyses and visualizations were all completed in R 4.1.0 (R Core Team, 2021). We separately compiled all eBird data recorded during summer (June-August) and nonbreeding season (December-February) for all species. For shorebirds (Charadriiformes), we limited summer months to June-July so their August movement would not affect distribution predictions, which are meant to approximate breeding ranges. In all checklists, we discarded subspecies information and summarized all data at the species level. We initially applied several filters to the data to reduce bias and improve data quality following established eBird data modeling protocols (Johnston et al., 2019; Kelling et al., 2018). First, we eliminated checklists covering extreme high durations (> 3 hours), with high numbers of observers (>5), or adhering to any protocol other than “stationary” or “traveling”, as these are incomparable with the bulk of eBird’s data. Second, to reduce positional error of observations, we eliminated checklists covering > 1 km because these are likely to result in greater spatial uncertainty. Third, we removed data before 2004 as there is too little data from earlier years to adequately control for long-term temporal trends. Finally, we removed data from any users with fewer than five contributions to reduce bias (e.g., false absence) resulting from inexperienced birders.

Data was further filtered at the species level before modeling. To limit overprediction outside of the species range extent, checklists were limited to those falling within a 200 km buffer of the species’ seasonal expert range boundary (via Cornell’s spatial boundaries - https://ebird.org/science/status-and-trends/download-data, accessed July 2021). We additionally excluded August data when modeling shorebirds (order Charadriiformes) as many species are already migrating long distances by this time. To reduce site-selection and temporal bias in data collection, we filtered checklists to one per 5 km grid cell per week. We repeated this filtering for checklists in which the species does and does not occur to reduce imbalance between presence and absence points. All coordinates, polygons and grids used in the study operated under a conical equal area projection. The *raster* (Hijmans et al., 2015), *rgdal* (Bivand et al., 2015), and *sf* (Pebesma, 2018) packages were used for spatial geoprocessing.

### Model covariates

Each SDM included several classes of covariates to account for numerous factors that influence species distributions. Our topographic/habitat suite of covariates included mean elevation (from EarthEnv; Robinson et al., 2014), mean enhanced vegetation index (EVI; MODIS; https://lpdaac.usgs.gov/products/mod11a1v006/), topographic wetness index (TWI; hydroSHEDS; Marthews et al., 2015), and terrain roughness index (TRI; EarthEnv). Climatic covariates included mean annual temperature (bio1), mean annual precipitation (bio12), precipitation seasonality (bio15; all bio variables from CHELSA v2.1; Karger et al., 2021), and intra-annual variation in cloud cover (EarthEnv). Covariates were selected from a pool of 32 contenders based on minimal collinearity. All covariates not available at 1 km were spatially aggregated to 1 km based on the mean value in each cell.

Our landcover suite of covariates included percent landcover within a buffer (with size corresponding to modeling grain size) corresponding to the year the checklist was recorded for the following categories: cropland, evergreen broadleaf, deciduous broadleaf, evergreen needleleaf, deciduous needleleaf, mixed forest, mosaic, shrubland, grassland, lichens/mosses, sparse, flooded/freshwater, flooded/saltwater, flooded/shrub, urban, barren, water, and ice (from European Space Agency – Climate Change Initiative; “ESA. Land Cover CCI Product User Guide Version 2. Technical Report.,” 2017). We also included temporal covariates – year, date, and time of day – in models to account for temporal variability in bird activity and long-term population trends. Finally, we included all effort indicators as covariates.

### Prediction surface

We initially generated a prediction surface covering our North American bounding box containing 1 km cells. We generated 1 km resolution values of all environmental covariates (topographic, climatic, and landcover suites) corresponding to each cell. When predicting the distribution of each species, we masked portions of the prediction surface that fell outside of the buffered range extent. We generated predictions for non-spatial covariates assuming a 1 km, 1 hour search with 1 observer in 2020 at a randomized date within the season and during the hour of the day when the species is observed at the highest rate.

### Spatial grain

To generate SDMs across spatial grains, we mean-aggregated all spatial covariates assigned to each point (topographic, climatic, and percent landcover suites) and our prediction surface to the following spatial grain sizes (test grains) from 1 km: 3 km, 5 km, 10 km, and 50 km. Test grains were selected based on their common usage in published SDMs. Temporal and effort covariates, which are assigned at the checklist (point) level, were not aggregated.

### Species distribution models

We separately modeled the summer and winter distributions of each species using Random Forest, a machine learning method designed to analyze large datasets with many covariates which is often found to generate the most accurate SDMs (e.g., Mi et al., 2017). Random forests are flexible, adjust automatically to complex, nonlinear relationships, and consider high-order interactions between predictors (Evans et al., 2011).

Before modeling, we randomly split data 60/20/20 into training, testing and out-of-bag (OOB) samples. We used the testing set for model validation and the OOB set for threshold estimation. Using the *ranger* package (Wright et al., 2018), we fit five models for each species during each season corresponding to each test grain, resulting in 4,955 separate models. Models were parameterized to 100 trees and 7 threads. We compiled model predictive performance metrics, including area under the curve (AUC), Kappa, true skill statistic (TSS), sensitivity, and specificity. We selected optimal thresholds for each model by maximizing the sum of sensitivity and specificity. Following each model fit, we generated thresholded grid-level occurrence predictions by predicting to the prediction surface. We further compiled the predicted area of occurrence and variable importance scores for covariates. For each model, we checked test-set calibration plots to diagnose overfitting.

### Aggregated biodiversity

We aggregated thresholded predictions of species distributions to produce species richness and range-size rarity (a measure of richness weighted towards species with smaller range sizes) for both seasons at every test grain. To generate aggregated richness, we summed the predicted occurrences for all species within every cell in our prediction surface. To generate range-size rarity, we divided the contribution of each species to the richness of a cell by its range size in km. Biodiversity visualizations were generated using color palettes from *Rcolorbrewer* (Neuwirth and Neuwirth, 2011).

### Aggregated Prediction Validation

To validate the aggregated richness predictions at each test grain, we compared model predictions to species count totals at 322 specific locations throughout the US and Canada. To ensure even spatial coverage of these locations, we selected the most heavily birded eBird location (minimum 100 visits per season, to ensure completeness of sightings) within every 200 km grid cell and compiled the total number of species and identities of all species recorded at that location for each season. We then compared hotspot richness and species identities against the aggregated richness predictions at each test grain and for each season. We calculated the percentage of individual species estimated to be present and absent that were identified correctly or incorrectly for the cell corresponding to each hotspot. To determine congruence in richness totals, we calculated the slopes between hotspot and estimated richness for all five test grains, as well as R^2^ values, root mean square error, and symmetric mean absolute percentage error (via the Metrics package; Hamner et al., 2018).

### Sensitivity across species and regions

To determine which species are most sensitive to modeling grain, we examined associations between the change in AUC (Δ AUC) between models fit at 1 km and 5 km grain sizes and several species-level factors: 1) landcover diversity index (LDI), a measure of habitat specialization, based on the heterogeneity in landcover covariate importance scores in each species’ seasonal random forest model (at 1 km; see Zuckerberg et al., 2016); 2) estimated range size at 1 km; and 3) the dominant biome or ecoregion (via Dinerstein et al., 2017) to which the species belongs. We square-root transformed range size prior to use in models. Ecoregions were categorically assigned at the species and seasonal level if >50% of the species range boundary was within one ecoregion type; otherwise, the species was designated as occupying “multiple” ecoregions. We manually corrected several ecoregion assignments among shorebirds (Charadriiformes), where purely coastal species were assigned adjacent inland ecoregions.

Using a multiple regression generalized linear model (GLM), we simultaneously modeled the influence of all three factors, plus season as a covariate, on the species-level Δ AUC between 1 km and 5 km grain sizes.

## Results

We fit and thresholded SDMs at five spatial grains for all bird species of the United States and Canada across two annual timepoints, amounting to 572 species during summer and 419 species during winter. Predicted ranges were highly dependent on grain size; for example, relative to fine-grain models, white-breasted nuthatch (*Sitta carolinensis*) lost significant pieces of its heterogenous western range in coarse-grain models but retained its homogenous eastern range (Fig 2a). Sedge wren (*Cistothorus stellaris*), which breeds in the prairie pothole region, and lesser goldfinch (*Spinus psaltria*), which breeds in southwestern deserts, each experienced deterioration of the range edge in coarse-grain models, while unsuitable areas closer to the range interior under fine-grain models became suitable under coarse models (Fig. 2b-c). Relative to 1 km models, species experienced both range loss and range gain when modeled at coarse grains, though range loss occurred more often (83% of species in summer and 63% in winter in 3 km vs. 1 km models). In summer, the median species lost ∼40% of its range at coarse grains, though median range loss was only ∼10% in winter (Fig. 2d). In cross-validation, model performance was high at all spatial grains for most species but was especially high at 1 km based on either AUC or TSS (Fig. 3).

**Figure 2.**
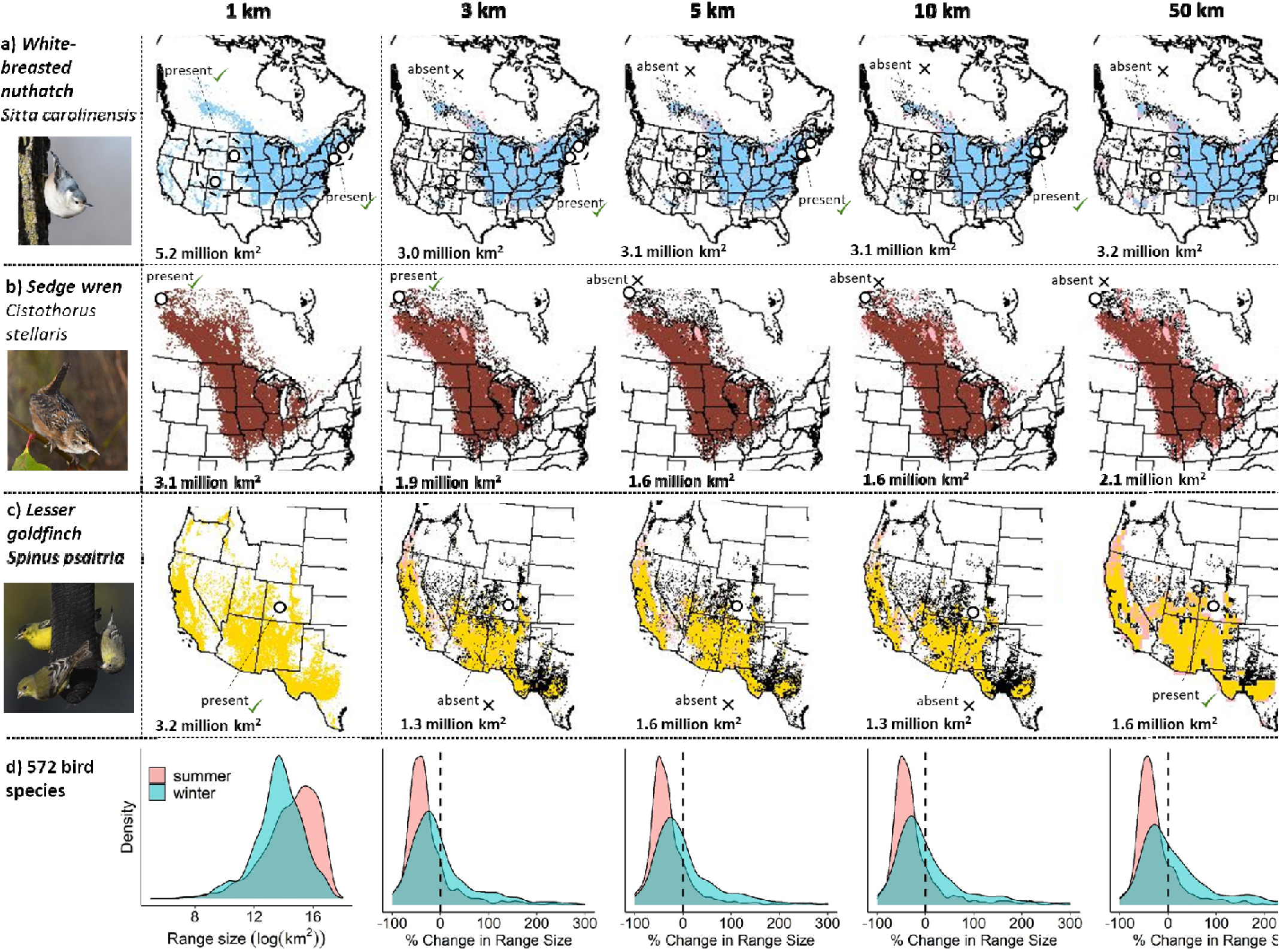
Dependence of species distribution predictions on grain size. Predicted breeding range of (a) white-breasted nuthatch (*Sitta carolinensis*), (b) sedge wren (*Cistothorus stellaris*), and (c) lesser goldfinch (*Spinus psaltria*) across five spatial modeling grains ranging from 1 km to 50 km. **For grains coarser than 1 km, colored areas represent predicted suitable area in both 1 km and coarser-grain models, black areas are suitable in 1 km but not coarser models, and pink areas are suitable in coarser-grain but not 1 km models.** Points correspond to five focal locations and are labeled with model-estimated presence or absence for the species and whether that estimation was correct based on validations. Locations were chosen based on their independent sizes and habitat types (see Fig. 3). Range size estimates for each model appear in bottom right corners. (d) Density plots show estimated area of occurrence for species distribution models fit at 1 km and change in area of occurrence for models fit at 3, 5, 10, and 50 km grain relative to 1 km models. Distributions are given for modeled summer (red) and winter (blue) distributions.

**Figure 3:**
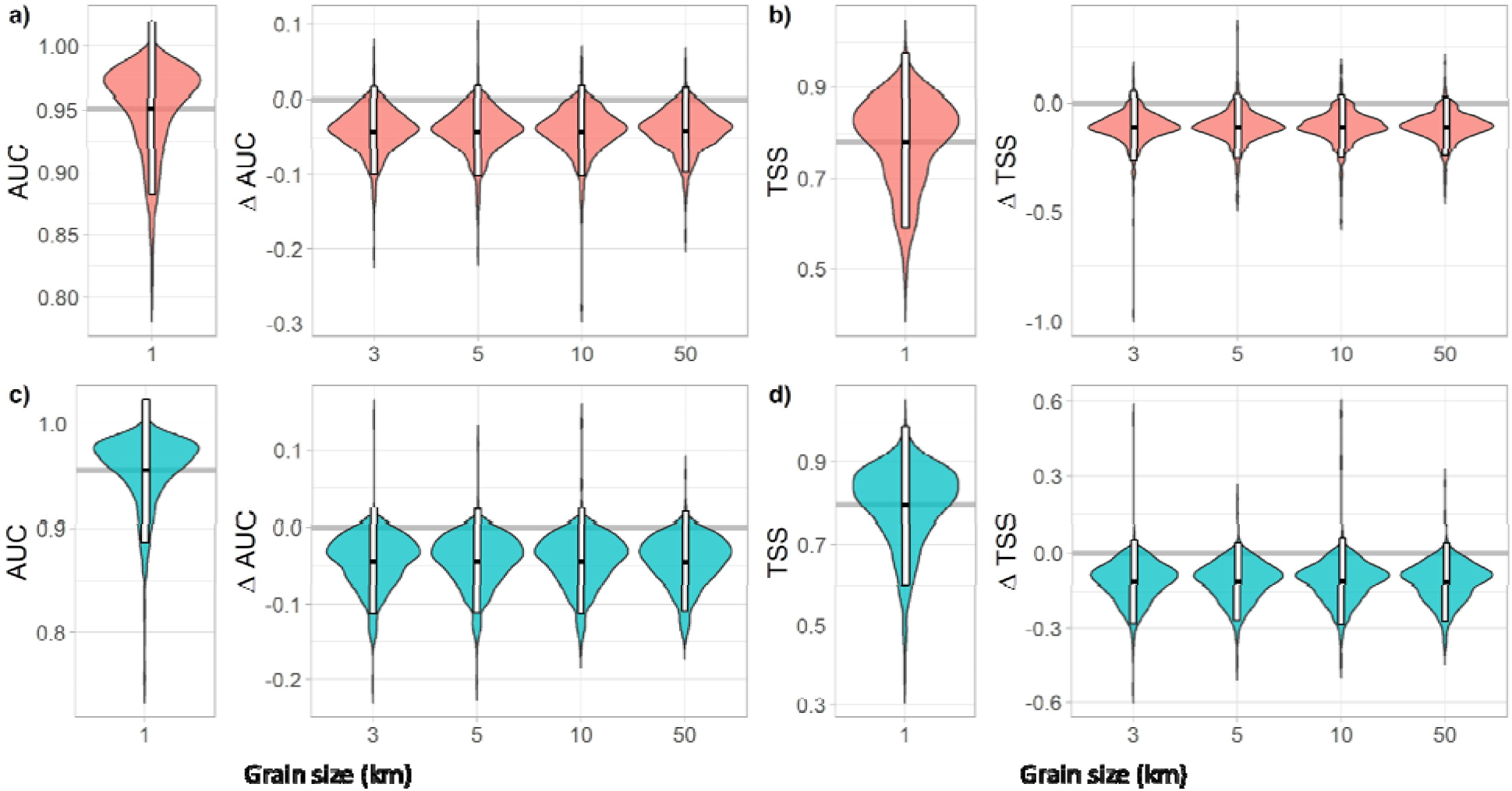
Model performance across spatial grains. Performance of species distribution models at 1 km across species (a,c, area under the curve; b,d, true skill statistic) and change in model performance between models fit at 3, 5, 10, and 50 km compared with 1 km after within-model cross-validation. Species distributions were modeled for (a-b, red) summer and (c-d, blue) winter. White bars give 95% intervals and black lines are medians.

Aggregating these maps reveals well-known regional differences in species richness (Fig. 4a-b, left; Fig. S1) and range-size rarity (Fig. S2, left) patterns. However, patterns differ substantially among predictions generated at different spatial grains. In summer, as spatial grain increased, models consistently overpredicted richness and range-size rarity in the American west and British Columbia, while underpredicting both in the boreal forest, prairie pothole region and American great plains (Fig 2a, right panels). In winter, models likewise overpredicted richness in the American west, though we did not observe underprediction elsewhere (Fig 2b, right panels). Predictions in the eastern deciduous forest, a region with more homogenous landcover, were highly consistent regardless of spatial grain.

**Figure 4.**
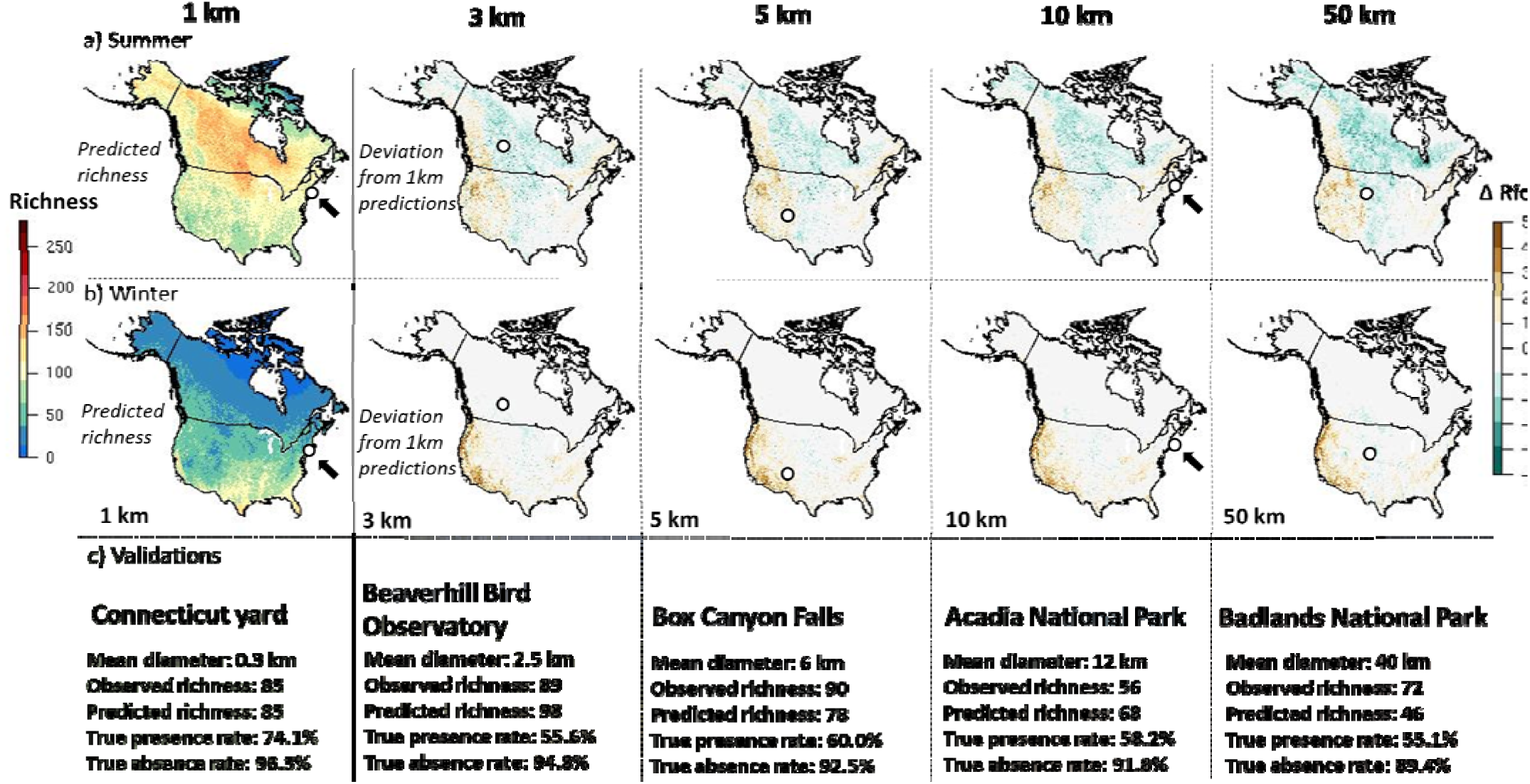
Grain-size dependence of avian species richness predictions. Left panels show predicted richness based on models trained, and predicted, at 1km grain for (a) summer and (b) winter. Left to right, panels show the difference between predicted richness based on models trained at successively coarser grain compared with predicted richness at 1 km. (c) Predictions of species richness and presence or absence were validated via comparisons to observations at five focal locations, chosen for their approximate correspondence in size to each model grain (points in a-b).

We validated predictions of richness and species presence and absence by assessing the accuracy of aggregated predictions across models through comparisons with site-level bird checklists. Comparisons with five focal birding hotspots, chosen because their approximate sizes correspond to our modeling grains, revealed variation in accuracy based on model grain (Fig. 4c). For example, observations in a large Connecticut, USA yard (diameter 0.3 km) closely matched richness predictions at 1 km, with a 74% true presence rate and 96% true absence rate. Meanwhile, observations at Badlands National Park (diameter 40 km) poorly matched predictions at 50 km, with a 55% true presence and 89% true absence rate. Likewise, when predicting the presence or absence of species at 322 locations, 1 km models outperformed coarser grain models, correctly predicting 68% of species present and 96% of species absent compared with rates of 55-60% and 94-95% for coarser models (Fig. 5). Similarly, coarse grain models incorrectly predicted species present or absent more often than 1 km models (Fig. S3). 1 km models also predicted total richness across sites more accurately than coarser models (Table S1; Fig. S4). These patterns were roughly consistent across seasons.

**Figure 5.**
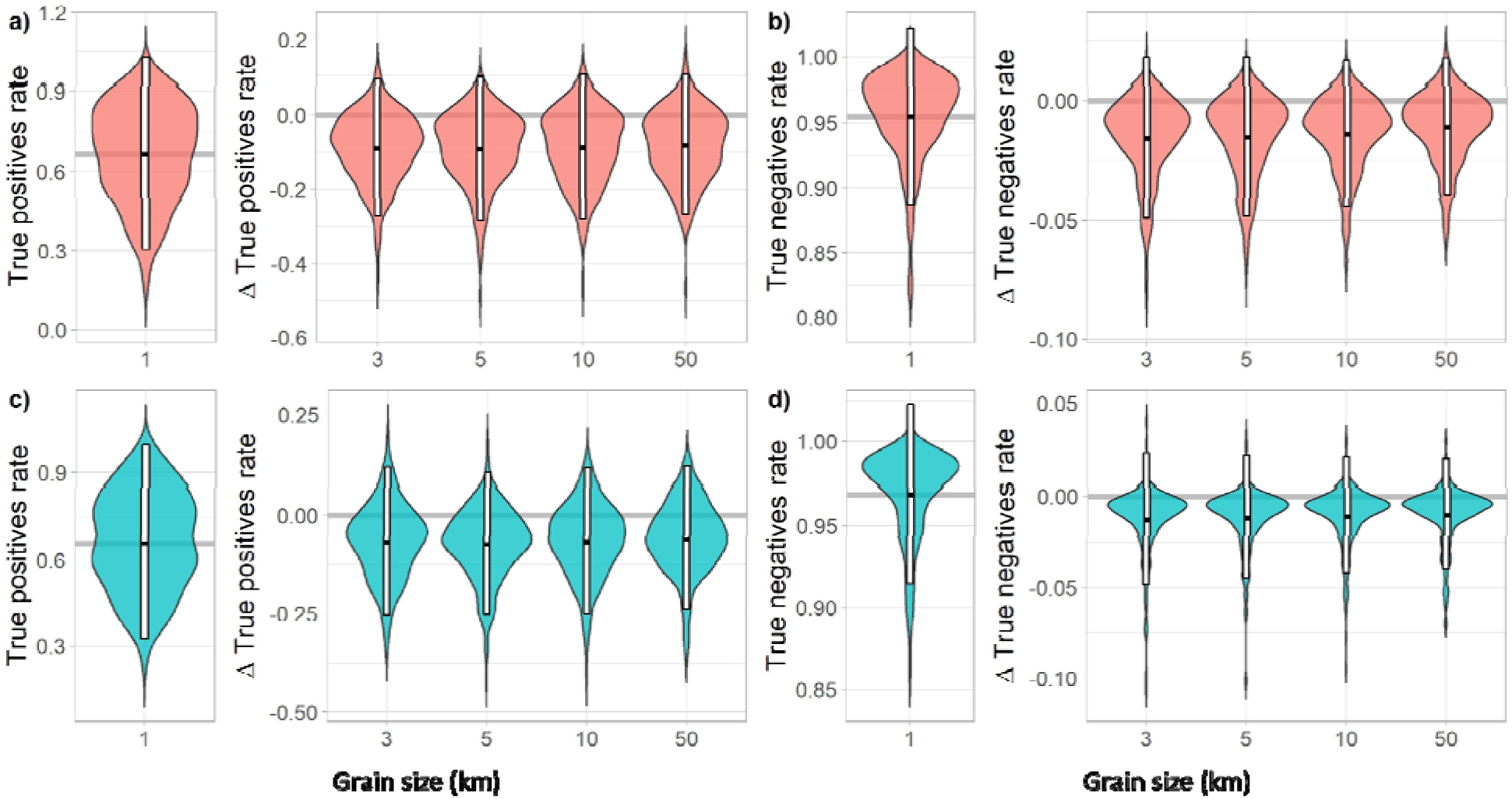
Performance of aggregated model predictions. Proportion of species correctly identified as present or absent at a set of 250 locations across the US and Canada for 1 km species distribution models (a,c, area under the curve; b,d, true skill statistic) and change in model performance between models fit at 3, 5, 10, and 50 km compared with 1 km models (see Fig. S3 for false positive/negative rates). Species distributions were modeled for (a-b, red) summer and (c-d, blue) winter. White bars give 95% intervals and black lines are medians.

Across species, we did not find that either habitat specialization (LDI) or range size was an important driver of the extent to which grain size impacted model performance as Δ AUC (generalized linear model and ANOVA: F<0.5, p>0.1; Table S2). However, the importance of model grain varied significantly by biome (F=8.1, p<0.001; Fig. 6; Table S2). Across both seasons, the deserts and xeric scrublands biome exhibited the greatest ΔAUC between models conducted at all coarser grains (contrasts between 1 km and 5 km displayed in Fig. 6). The tundra, boreal forests and taiga, and tropical and subtropical moist broadleaf forests were less sensitive to grain size.

**Figure 6.**
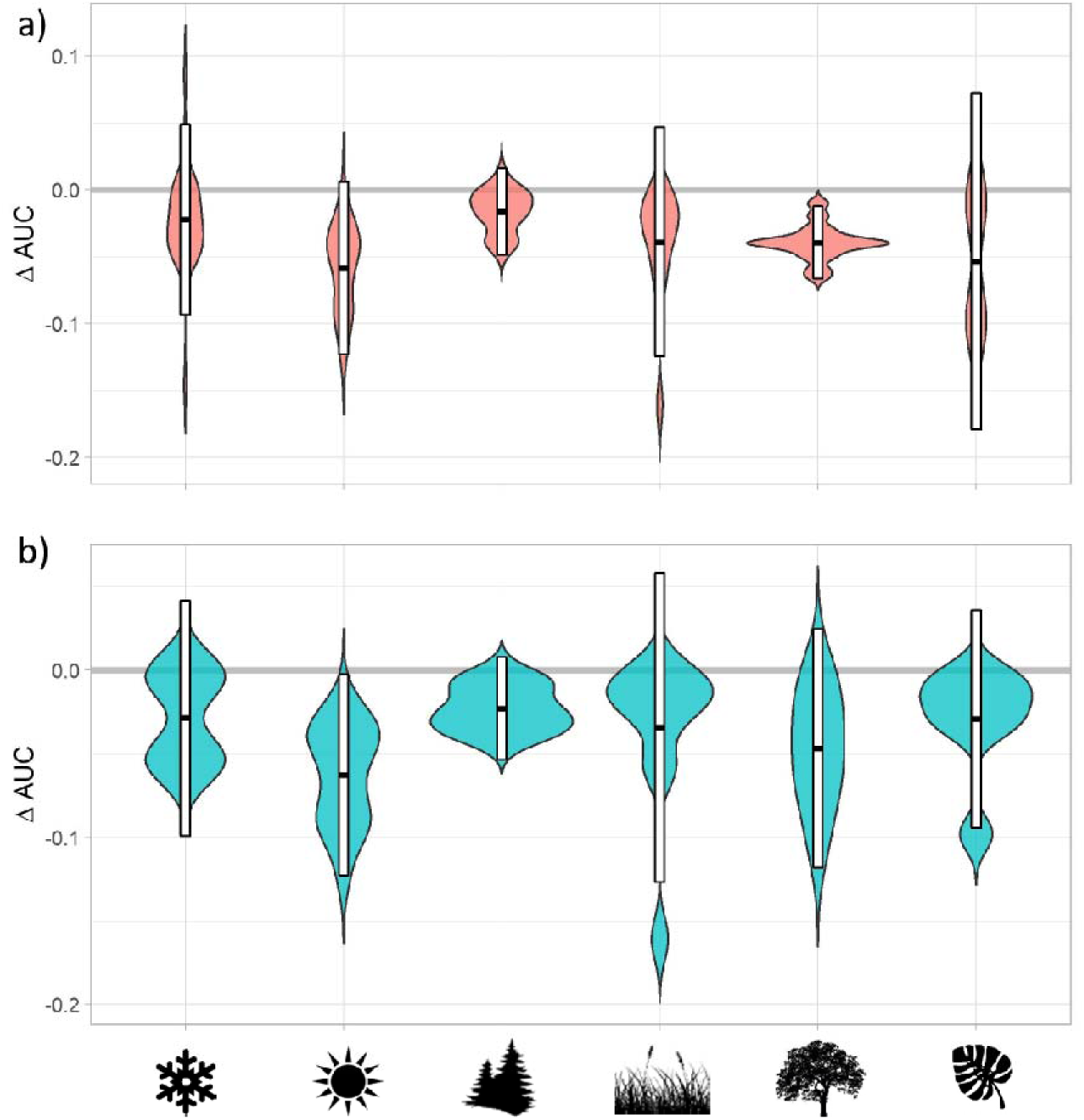
Biases related to grain size vary in intensity by habitat type. Change in species distribution model performance (Δ area under the curve) between models fit at 1 km vs. 5 km grain size compared with each species’ primary biome for (a) summer and (b) winter distributions. Silhouettes represent biomes, from left to right: tundra, deserts & xeric scrublands, boreal forests/ taiga, temperate grasslands and shrublands, temperate broadleaf and mixed forests, and tropical and subtropical moist broadleaf forests. White bars give 95% intervals and black lines are medians.

## Discussion

Modeling across spatial grain sizes is an important step towards understanding the biases present in existing coarse-grain estimates of continental species distributions and biodiversity patterns (Chauvier et al., 2022), widely used in climate change modeling and applied conservation efforts. While most cross-grain studies have concluded that fine-grain models are more reliable than coarse models (Seo et al., 2009; Song et al., 2013; Turner et al., 2019), some have suggested that grain size makes little difference (Guisan et al., 2007; Manzoor et al., 2018). Sometimes, modeling at coarser grains is encouraged, especially when certainty about the accuracy of the observation locations is poor (Gábor et al., 2020). Here, we evaluated the importance of spatial grain by modeling the continental distributions of 572 North American bird species at grain sizes of 1 to 50 km across summer and winter seasons and predicting biodiversity patterns. We cross-validated each model with independent test set data and validated presence, absence, and richness predictions using checklists at local sites. Our results suggest that model predictions at 1km are significantly more accurate than those fit at coarser grains, even at 3km. Thus, our results suggest that the increased accuracy of environmental covariate information at fine grains likely overcomes increasing uncertainty about the accuracy of observation locations (Gábor et al., 2022; Šímová et al., 2019), with the important caveat that we eliminated the most spatially-inaccurate observations (with observers covering distances > 1 km) from our dataset.

It was unclear whether coarse-grain models would systematically under- or over-estimate range sizes (thus similarly biasing richness when distributions are aggregated). Coarse-grain models may underestimate range sizes when large cells are considered unsuitable for a species despite local presence of suitable habitat or microclimate but may overestimate range sizes when entire cells are regularly considered suitable (Oldfather et al., 2020). We find that species distribution models most often underestimated, rather than overestimated range sizes, with even models at 3 km grain underestimating range area for 83% of species during summer and 63% of species during winter. Given this bias, many existing predictions of continental and global biodiversity conducted at coarse spatial grains (e.g., Mi et al., 2023; Stewart et al., 2022) may be overestimating richness and failing to characterize the full extent of biodiversity declines, especially as species are restricted to smaller habitat fragments (Bergerot et al., 2012). This bias may also be a function of whether models are presence-absence, as they are here, or presence-only. Presence-only models have less precise absence information, especially when species are under-sampled, and may associate more unsuitable habitat or climate space with presence points (Fiedler et al., 2018); thus, we encourage further exploration of bias in predicted range sizes using presence-only models.

Biases resulting from modeling at coarse grains may predictably intensify for certain regions or species and during specific times of year (Mateo Sánchez et al., 2014), creating important seasonal and regional biases in biodiversity predictions when aggregated. Our models and biodiversity aggregations reveal that while grain-related bias is present during both seasons, grain size most consistently biased area-of-occurrence estimates during summer, despite similar decreases in model performance based on cross-validation across seasons. We find that regions with highly heterogenous landscapes, such as the American southwest, are likely to be most prone to misestimation of area of occurrence, while models in homogenous regions, e.g. the boreal and taiga biomes and the American southeast, may be less affected. Meanwhile, biases in richness predictions occurred in both directions; coarse-grain models overpredicted diversity in the west and underpredicted it in the great plains, prairie pothole region and boreal/taiga zones. Finally, while we expected that species with small ranges or high habitat specificity may be especially prone to loss of estimated range area at coarse grains, we did not detect an effect of these attributes on grain-related bias. Thus, we find that grain-related bias is most pronounced across different regions and seasons but not species.

Although the finest grain size in our models was 1 km, species distribution models are possible at even finer resolutions, including 0.5 km or below (Manzoor et al., 2018), given the increasing availability of microclimate and microhabitat information (He et al., 2015; Urbina-Cardona et al., 2019). Models fit at 0.5 km may be more accurate than those at 1 km if observational and environmental data is very precise, spatial extents are small, and home ranges are highly localized (Connor et al., 2019, 2018). However, several limitations preclude the feasibility of 0.5 km or smaller-grain SDMs at continental scales. For example, across continental extents, computational limitations may become severe when modeling at 0.5 km with over 100 million cells in North America alone. Even at sub-continental extents, very high-resolution models become less accurate with increasing extent (Connor et al., 2019). Many weather and topographic products are still unavailable below 1km, especially at high temporal resolution, and would require interpolation and reductions in temporal scale to achieve that resolution. Further, the operational grain of many bird and mammal species is greater than 0.5 km, even during summer when species are breeding and highly localized (Mertes and Jetz, 2018), although sub-1km grain models might perform better for more stationary organisms such as plants, corals, insects, or reptiles. Finally, the positional uncertainty of observations is likely to become especially important at grains below 1 km (Gábor et al., 2020) as large numbers of observations (27% of our checklists) cover at least half a kilometer of distance. These reasons may explain why ultra-fine grain SDMs have not always produced more accurate results (Manzoor et al., 2018).

### Conclusions

To understand changing biodiversity patterns at both local and regional scales, researchers, conservationists, and managers rely on accurate predictions of species distributions and their environmental drivers (Araújo et al., 2019; Jetz et al., 2008). Modeling distributional patterns across spatial grains is an important step towards understand the biases present in existing estimates that are widely used in climate change modeling and applied conservation efforts (Chauvier et al., 2022; Mertes and Jetz, 2018). Here, we show that when modeling the distributions of 572 bird species across two seasons, fine-grain SDMs provide more realistic assessments of species distributions, predicted area of occurrence, and biodiversity. Researchers must be cognizant of spatial grain and its influence on our understanding of species distributions and biodiversity to ensure the best possible conservation outcomes in the face of global change.

## Supplementary Information

**Table S1.** Slope coefficient, R^2^ value and error (rmse = root mean square error; smape = symmetric mean absolute percentage error) based on independent validation of richness predictions across five spatial grain sizes and two seasons.

| Season | Scale (km) | Slope | $R^2$ | rmse | smape |
| --- | --- | --- | --- | --- | --- |
| Summer | 1 | 0.262 | 0.182 | 32.529 | 0.325 |
|  | 3 | 0.082 | 0.043 | 35.442 | 0.363 |
|  | 5 | 0.074 | 0.036 | 35.492 | 0.363 |
|  | 10 | 0.085 | 0.046 | 35.089 | 0.361 |
|  | 50 | 0.098 | 0.069 | 34.212 | 0.360 |
| Winter | 1 | 0.555 | 0.687 | 22.162 | 0.285 |
|  | 3 | 0.453 | 0.421 | 29.648 | 0.355 |
|  | 5 | 0.439 | 0.446 | 28.759 | 0.348 |
|  | 10 | 0.460 | 0.459 | 28.303 | 0.344 |
|  | 50 | 0.421 | 0.433 | 28.994 | 0.348 |

**Table S2.** ANOVA table summarizing associations between species-level attributes and change in AUC between species distribution models fit at 1 km and 5 km grain sizes. ANOVA was fit to a generalized linear model.

| ANOVA table | Sums of Squares | DF | F-value | p-value |
| --- | --- | --- | --- | --- |
| Landcover diversity Index | <0.001 | 1.000 | 0.045 | 0.832 |
| Range size | <0.001 | 1.000 | 0.463 | 0.496 |
| Primary biome | 0.042 | 6.000 | 8.088 | <0.001 |
| Season | 0.004 | 2.000 | 2.304 | 0.100 |
| Residuals | 0.814 | 946.000 |  |  |

**Figure S1.**
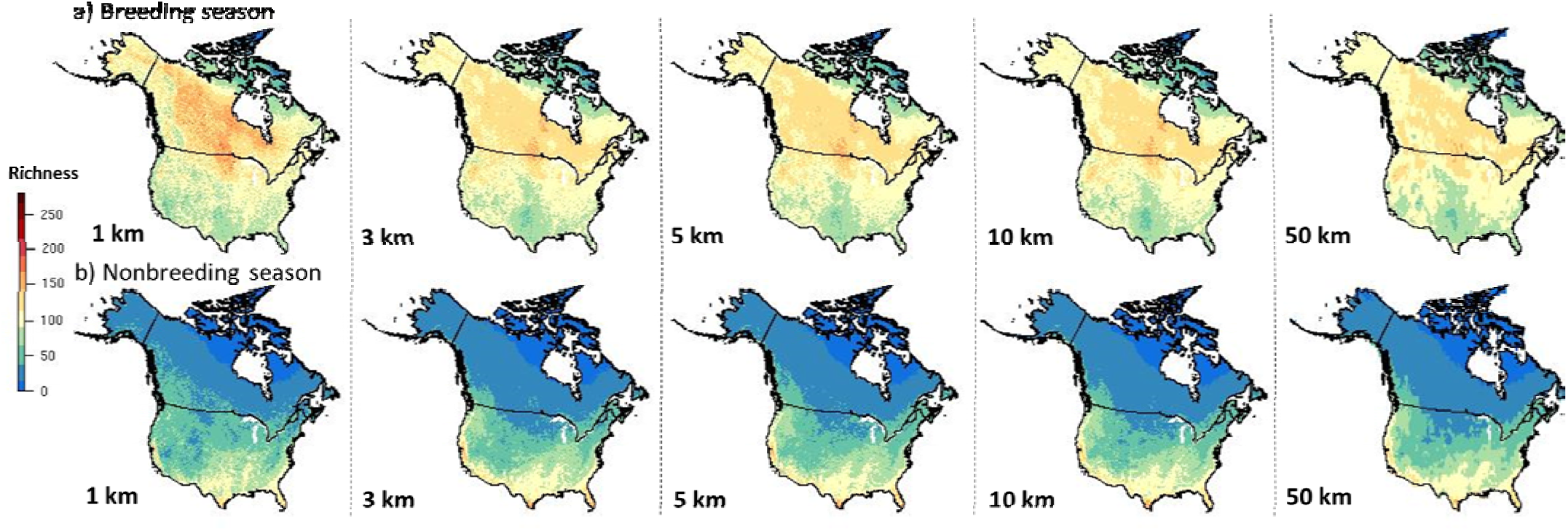
Estimated biodiversity across scales. Estimated continental avian richness at 1 km grain size in the United States and Canada for (a) summer and (b) winter.

**Figure S2.**
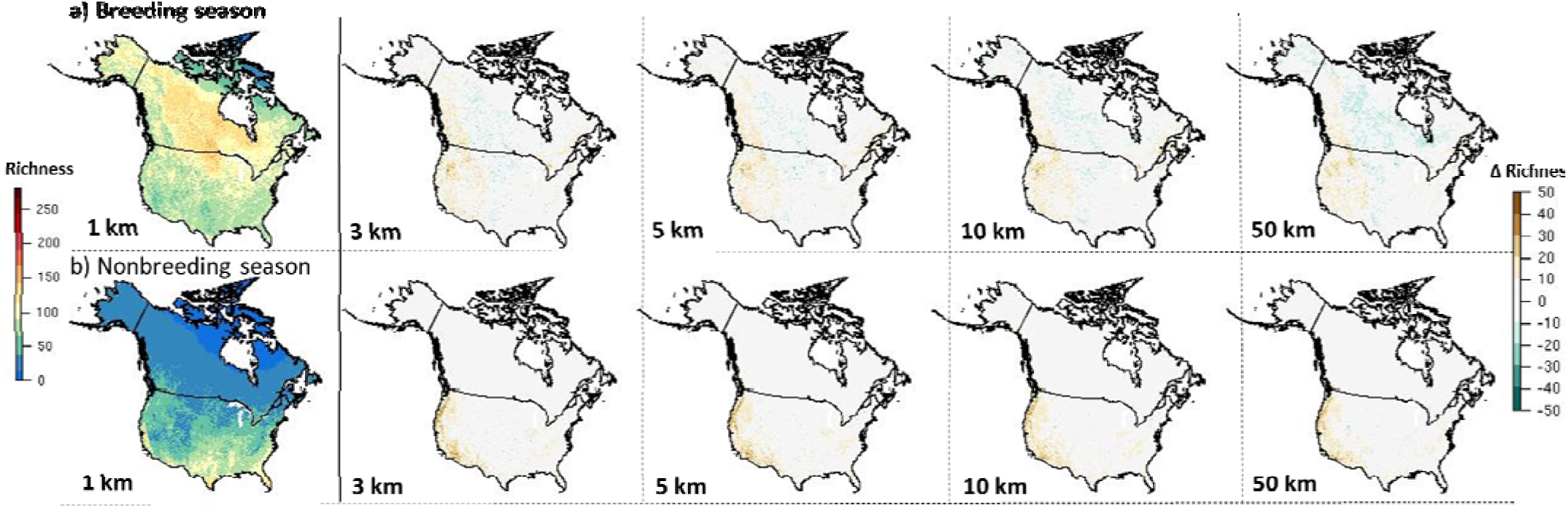
Delta range-size rarity across scales. Estimated continental avian range-size rarity at 1 km grain size in the United States and Canada (left panels) and difference in range-size rarity between 1 km and other grain sizes (right panels) for (a) summer and (b) winter.

**Figure S3.**
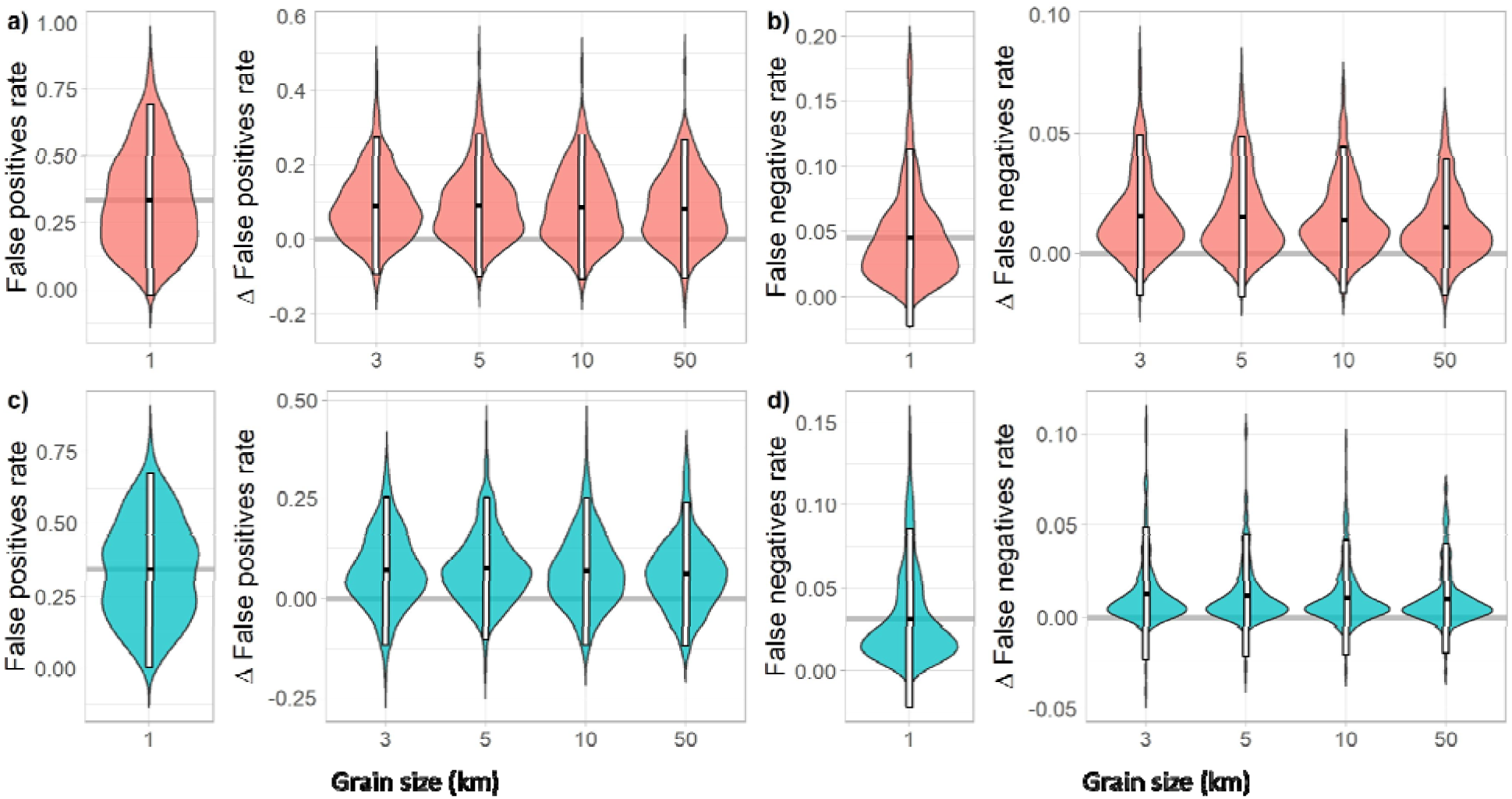
Proportion of species incorrectly identified as present or absent at a set of 250 locations across the US and Canada for 1 km models (a,c, area under the curve; b,d, true skill statistic) and change in model performance between models fit at 3, 5, 10, and 50 km compared with 1 km models. Species distributions were modeled for (a-b, red) summer and (c-d, blue) winter. White bars give 95% intervals and black lines are medians.

**Figure S4.**
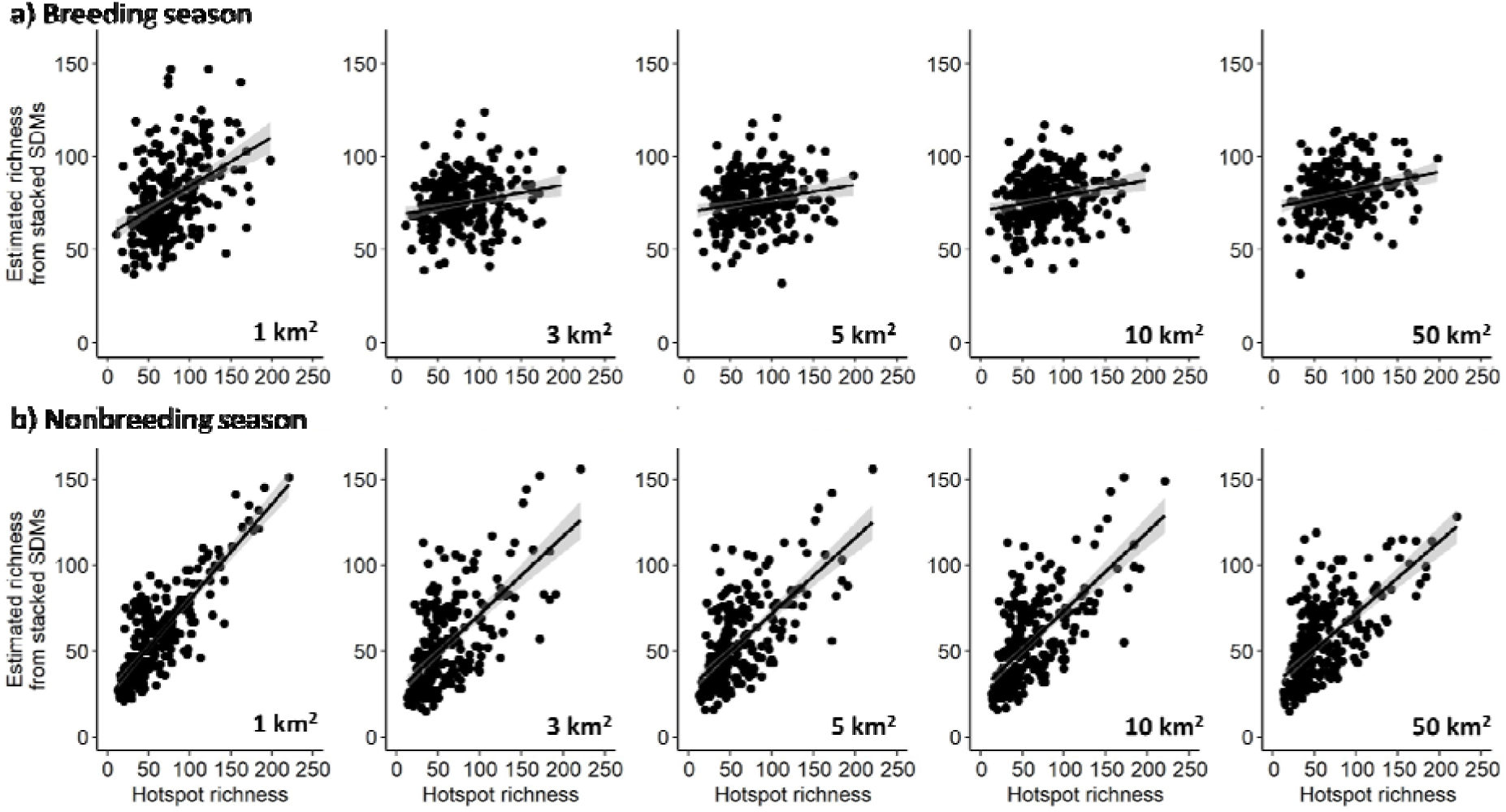
Accuracy of richness predictions generated based on species distribution models across five spatial grains and two seasons, (a) summer and (b) winter.

## Notes

### Competing Interest Statement

The authors have declared no competing interest.

## References

A. Lee-Yaw, J., L. McCune, J., Pironon, S., N. Sheth, S., 2022. Species distribution models rarely predict the biology of real populations. Ecography 2022, e05877.

Abrahms, B., DiPietro, D., Graffis, A., Hollander, A., 2017. Managing biodiversity under climate change: challenges, frameworks, and tools for adaptation. Biodivers. Conserv. 26, 2277–2293. 10.1007/s10531-017-1362-4

Araújo, M.B., Anderson, R.P., Márcia Barbosa, A., Beale, C.M., Dormann, C.F., Early, R., Garcia, R.A., Guisan, A., Maiorano, L., Naimi, B., 2019. Standards for distribution models in biodiversity assessments. Sci. Adv. 5, eaat4858.

Association, A.B., 2008. American Birding Association Checklist: Birds of the Continental United States and Canada.

Austin, M.P., Van Niel, K.P., 2011. Improving species distribution models for climate change studies: variable selection and scale: Species distribution models for climate change studies. J. Biogeogr. 38, 1–8. 10.1111/j.1365-2699.2010.02416.x

Barbet-Massin, M., Jetz, W., 2015. The effect of range changes on the functional turnover, structure and diversity of bird assemblages under future climate scenarios. Glob. Change Biol. 21, 2917–2928. 10.1111/gcb.12905

Beery, S., Cole, E., Parker, J., Perona, P., Winner, K., 2021. Species distribution modeling for machine learning practitioners: a review, in: ACM SIGCAS Conference on Computing and Sustainable Societies. pp. 329–348.

Bergerot, B., Merckx, T., Van Dyck, H., Baguette, M., 2012. Habitat fragmentation impacts mobility in a common and widespread woodland butterfly: do sexes respond differently? BMC Ecol. 12, 5. 10.1186/1472-6785-12-5

Bivand, R., Keitt, T., Rowlingson, B., Pebesma, E., Sumner, M., Hijmans, R., Rouault, E., Bivand, M.R., 2015. Package ‘rgdal.’ Bind. Geospatial Data Abstr. Libr. Available Online Httpscran R-Proj. Orgwebpackagesrgdalindex Html Accessed 15 Oct. 2017.

Buckley, L.B., Jetz, W., 2008. Linking global turnover of species and environments. Proc. Natl. Acad. Sci. 105, 17836–17841. 10.1073/pnas.0803524105

Carroll, K.A., Farwell, L.S., Pidgeon, A.M., Razenkova, E., Gudex-Cross, D., Helmers, D.P., Lewińska, K.E., Elsen, P.R., Radeloff, V.C., 2022. Mapping breeding bird species richness at management-relevant resolutions across the United States. Ecol. Appl. 32, e2624.

Chauvier, Y., Descombes, P., Guéguen, M., Boulangeat, L., Thuiller, W., Zimmermann, N.E., 2022. Resolution in species distribution models shapes spatial patterns of plant multifaceted diversity. Ecography 2022, e05973.

Clavel, J., Julliard, R., Devictor, V., 2011. Worldwide decline of specialist species: toward a global functional homogenization? Front. Ecol. Environ. 9, 222–228.

Clements, J.F., 2007. Clements checklist of birds of the world. Comstock Pub. Associates/Cornell University Press.

Connor, T., Hull, V., Viña, A., Shortridge, A., Tang, Y., Zhang, J., Wang, F., Liu, J., 2018. Effects of grain size and niche breadth on species distribution modeling. Ecography 41, 1270– 1282. 10.1111/ecog.03416

Connor, T., Viña, A., Winkler, J.A., Hull, V., Tang, Y., Shortridge, A., Yang, H., Zhao, Z., Wang, F., Zhang, J., Zhang, Z., Zhou, C., Bai, W., Liu, J., 2019. Interactive spatial scale effects on species distribution modeling: The case of the giant panda. Sci. Rep. 9, 14563. 10.1038/s41598-019-50953-z

Dinerstein, E., Olson, D., Joshi, A., Vynne, C., Burgess, N.D., Wikramanayake, E., Hahn, N., Palminteri, S., Hedao, P., Noss, R., 2017. An ecoregion-based approach to protecting half the terrestrial realm. BioScience 67, 534–545.

Distler, T., Schuetz, J.G., Velásquez-Tibatá, J., Langham, G.M., 2015. Stacked species distribution models and macroecological models provide congruent projections of avian species richness under climate change. J. Biogeogr. 42, 976–988. 10.1111/jbi.12479

ESA. Land Cover CCI Product User Guide Version 2. Technical Report., 2017.

Evans, J.S., Murphy, M.A., Holden, Z.A., Cushman, S.A., 2011. Modeling species distribution and change using random forest, in: Predictive Species and Habitat Modeling in Landscape Ecology. Springer, pp. 139–159.

Fiedler, P.C., Redfern, J.V., Forney, K.A., Palacios, D.M., Sheredy, C., Rasmussen, K., García-Godos, I., Santillán, L., Tetley, M.J., Félix, F., 2018. Prediction of large whale distributions: a comparison of presence–absence and presence-only modeling techniques. Front. Mar. Sci. 419.

Gábor, L., Jetz, W., Lu, M., Rocchini, D., Cord, A., Malavasi, M., Zarzo-Arias, A., Barták, V., Moudrỳ, V., 2022. Positional errors in species distribution modelling are not overcome by the coarser grains of analysis. Methods Ecol. Evol. 13, 2289–2302.

Gábor, L., Moudrỳ, V., Lecours, V., Malavasi, M., Barták, V., Fogl, M., Šímová, P., Rocchini, D., Václavík, T., 2020. The effect of positional error on fine scale species distribution models increases for specialist species. Ecography 43, 256–269.

Gaiji, S., Chavan, V., Ariño, A.H., Otegui, J., Hobern, D., Sood, R., Robles, E., 2013. Content assessment of the primary biodiversity data published through GBIF network: status, challenges and potentials. Biodivers. Inform. 8.

Gillingham, P.K., Huntley, B., Kunin, W.E., Thomas, C.D., 2012. The effect of spatial resolution on projected responses to climate warming. Divers. Distrib. 18, 990–1000. 10.1111/j.1472-4642.2012.00933.x

Gottschalk, T.K., Aue, B., Hotes, S., Ekschmitt, K., 2011. Influence of grain size on species– habitat models. Ecol. Model. 222, 3403–3412. 10.1016/j.ecolmodel.2011.07.008

Guisan, A., Graham, C.H., Elith, J., Huettmann, F., the NCEAS Species Distribution Modelling Group, 2007. Sensitivity of predictive species distribution models to change in grain size. Divers. Distrib. 13, 332–340. 10.1111/j.1472-4642.2007.00342.x

Hamner, B., Frasco, M., LeDell, E., 2018. Package ‘Metrics.’ R Found. Stat. Comput.

He, F., Condit, R., 2007. The distribution of species: occupancy, scale, and rarity. Scaling Biodivers. 32–50.

He, K.S., Bradley, B.A., Cord, A.F., Rocchini, D., Tuanmu, M.-N., Schmidtlein, S., Turner, W., Wegmann, M., Pettorelli, N., 2015. Will remote sensing shape the next generation of species distribution models? Remote Sens. Ecol. Conserv. 1, 4–18. 10.1002/rse2.7

Hijmans, R.J., Van Etten, J., Cheng, J., Mattiuzzi, M., Sumner, M., Greenberg, J.A., Lamigueiro, O.P., Bevan, A., Racine, E.B., Shortridge, A., 2015. Package ‘raster.’ R Package 734.

Hurlbert, A.H., Jetz, W., 2007. Species richness, hotspots, and the scale dependence of range maps in ecology and conservation. Proc. Natl. Acad. Sci. 104, 13384–13389. 10.1073/pnas.0704469104

Jetz, W., Sekercioglu, C.H., Watson, J.E.M., 2008. Ecological Correlates and Conservation Implications of Overestimating Species Geographic Ranges: *Overestimation of Species Ranges*. Conserv. Biol. 22, 110–119. 10.1111/j.1523-1739.2007.00847.x

Johnston, A., Hochachka, W., Strimas-Mackey, M., Gutierrez, V.R., Robinson, O., Miller, E., Auer, T., Kelling, S., Fink, D., 2019. Best practices for making reliable inferences from citizen science data: case study using eBird to estimate species distributions. 10.1101/574392

Jung, M., Arnell, A., De Lamo, X., García-Rangel, S., Lewis, M., Mark, J., Merow, C., Miles, L., Ondo, I., Pironon, S., 2021. Areas of global importance for conserving terrestrial biodiversity, carbon and water. Nat. Ecol. Evol. 5, 1499–1509.

Karger, D.N., Wilson, A.M., Mahony, C., Zimmermann, N.E., 2021. Global daily 1km land surface precipitation based on cloud cover-informed downscaling 20.

Kelling, S., Johnston, A., Fink, D., Ruiz-Gutierrez, V., Bonney, R., Bonn, A., Fernandez, M., Hochachka, W.M., Julliard, R., Kraemer, R., Guralnick, R., 2018. Finding the signal in the Noise of Citizen Science Observations (preprint). Ecology. 10.1101/326314

Lecours, V., Devillers, R., Schneider, D., Lucieer, V., Brown, C., Edinger, E., 2015. Spatial scale and geographic context in benthic habitat mapping: review and future directions. Mar. Ecol. Prog. Ser. 535, 259–284. 10.3354/meps11378

Levin, S.A., 1992. The problem of pattern and scale in ecology: the Robert H. MacArthur award lecture. Ecology 73, 1943–1967.

Lu, M., Jetz, W., 2023. Scale-sensitivity in the measurement and interpretation of environmental niches. Trends Ecol. Evol.

Manzoor, S.A., Griffiths, G., Lukac, M., 2018. Species distribution model transferability and model grain size – finer may not always be better. Sci. Rep. 8, 7168. 10.1038/s41598-018-25437-1

Marthews, T.R., Dadson, S.J., Lehner, B., Abele, S., Gedney, N., 2015. High-resolution global topographic index values for use in large-scale hydrological modelling. Hydrol. Earth Syst. Sci. 19, 91–104.

Mateo Sánchez, M.C., Cushman, S.A., Saura, S., 2014. Scale dependence in habitat selection: the case of the endangered brown bear (*Ursus arctos*) in the Cantabrian Range (NW Spain). Int. J. Geogr. Inf. Sci. 28, 1531–1546. 10.1080/13658816.2013.776684

Mertes, K., Jetz, W., 2018. Disentangling scale dependencies in species environmental niches and distributions. Ecography 41, 1604–1615. 10.1111/ecog.02871

Mi, C., Huettmann, F., Guo, Y., Han, X., Wen, L., 2017. Why choose Random Forest to predict rare species distribution with few samples in large undersampled areas? Three Asian crane species models provide supporting evidence. PeerJ 5, e2849.

Mi, C., Ma, L., Yang, M., Li, X., Meiri, S., Roll, U., Oskyrko, O., Pincheira-Donoso, D., Harvey, L.P., Jablonski, D., 2023. Global Protected Areas as refuges for amphibians and reptiles under climate change. Nat. Commun. 14, 1389.

Moudrý, V., Šímová, P., 2012. Influence of positional accuracy, sample size and scale on modelling species distributions: a review. Int. J. Geogr. Inf. Sci. 26, 2083–2095. 10.1080/13658816.2012.721553

Neuwirth, E., Neuwirth, M.E., 2011. Package ‘RColorBrewer.’ CRAN 2011–06-17 08: 34: 00. Apache License 2.0.

Oldfather, M.F., Kling, M.M., Sheth, S.N., Emery, N.C., Ackerly, D.D., 2020. Range edges in heterogeneous landscapes: Integrating geographic scale and climate complexity into range dynamics. Glob. Change Biol. 26, 1055–1067.

Pacifici, M., Foden, W.B., Visconti, P., Watson, J.E.M., Butchart, S.H.M., Kovacs, K.M., Scheffers, B.R., Hole, D.G., Martin, T.G., Akçakaya, H.R., Corlett, R.T., Huntley, B., Bickford, D., Carr, J.A., Hoffmann, A.A., Midgley, G.F., Pearce-Kelly, P., Pearson, R.G., Williams, S.E., Willis, S.G., Young, B., Rondinini, C., 2015. Assessing species vulnerability to climate change. Nat. Clim. Change 5, 215–224. 10.1038/nclimate2448

Pebesma, E.J., 2018. Simple features for R: standardized support for spatial vector data. R J 10, 439.

R Core Team, 2021. R: A Language and Environment for Statistical Computing. R Foundation for Statistical Computing, Vienna, Austria.

Robinson, N., Regetz, J., Guralnick, R.P., 2014. EarthEnv-DEM90: A nearly-global, void-free, multi-scale smoothed, 90m digital elevation model from fused ASTER and SRTM data. ISPRS J. Photogramm. Remote Sens. 87, 57–67.

Seo, C., Thorne, J.H., Hannah, L., Thuiller, W., 2009. Scale effects in species distribution models: implications for conservation planning under climate change. Biol. Lett. 5, 39–43. 10.1098/rsbl.2008.0476

Šímová, P., Moudrý, V., Komárek, J., Hrach, K., Fortin, M.-J., 2019. Fine scale waterbody data improve prediction of waterbird occurrence despite coarse species data. Ecography 42, 511–520. 10.1111/ecog.03724

Song, W., Kim, E., Lee, D., Lee, M., Jeon, S.-W., 2013. The sensitivity of species distribution modeling to scale differences. Ecol. Model. 248, 113–118. 10.1016/j.ecolmodel.2012.09.012

Stewart, P.S., Voskamp, A., Santini, L., Biber, M.F., Devenish, A.J.M., Hof, C., Willis, S.G., Tobias, J.A., 2022. Global impacts of climate change on avian functional diversity. Ecol. Lett. 25, 673–685. 10.1111/ele.13830

Sullivan, B.L., Aycrigg, J.L., Barry, J.H., Bonney, R.E., Bruns, N., Cooper, C.B., Damoulas, T., Dhondt, A.A., Dietterich, T., Farnsworth, A., Fink, D., Fitzpatrick, J.W., Fredericks, T., Gerbracht, J., Gomes, C., Hochachka, W.M., Iliff, M.J., Lagoze, C., La Sorte, F.A., Merrifield, M., Morris, W., Phillips, T.B., Reynolds, M., Rodewald, A.D., Rosenberg, K.V., Trautmann, N.M., Wiggins, A., Winkler, D.W., Wong, W.-K., Wood, C.L., Yu, J., Kelling, S., 2014. The eBird enterprise: An integrated approach to development and application of citizen science. Biol. Conserv. 169, 31–40. 10.1016/j.biocon.2013.11.003

Thuiller, W., Guéguen, M., Renaud, J., Karger, D.N., Zimmermann, N.E., 2019. Uncertainty in ensembles of global biodiversity scenarios. Nat. Commun. 10, 1446.

Turner, J.A., Babcock, R.C., Kendrick, G.A., Hovey, R.K., 2019. How does spatial resolution affect model performance? A case for ensemble approaches for marine benthic mesophotic communities. J. Biogeogr. 46, 1249–1259. 10.1111/jbi.13581

Urbina-Cardona, N., Blair, M.E., Londoño, M.C., Loyola, R., Velásquez-Tibatá, J., Morales-Devia, H., 2019. Species distribution modeling in Latin America: a 25-year retrospective review. Trop. Conserv. Sci. 12, 1940082919854058.

Wiens, J.A., 1989. Spatial scaling in ecology. Funct. Ecol. 3, 385–397.

Wright, M.N., Wager, S., Probst, P., Wright, M.M.N., 2018. Package ‘ranger.’

Zuckerberg, B., Fink, D., La Sorte, F.A., Hochachka, W.M., Kelling, S., 2016. Novel seasonal land cover associations for eastern North American forest birds identified through dynamic species distribution modelling. Divers. Distrib. 22, 717–730. 10.1111/ddi.12428

